# flashscenic: Accelerated GRN Analysis with RegDiffusion for Large-Scale Single-Cell Data

**DOI:** 10.64898/2026.10.26.761795

**Authors:** Hao Zhu, Gonzalo Cardenal-Antolin, Xuting Jin, Jonathan S. Gootenberg, Omar O. Abudayyeh, Donna K. Slonim

**Author notes:** These authors contributed equally to this work (Hao Zhu, Gonzalo Cardenal-Antolin).

## Abstract

**Motivation:** Single-cell gene regulatory network (GRN) analysis via the SCENIC framework provides biologically interpretable cell-state representations through motif-validated transcription factor (TF) regulon activities. However, sequential, CPU-bound implementations (pySCENIC) constrain scalability to highly variable genes and make the repeated executions needed to resolve network stochasticity computationally prohibitive. Furthermore, while regulon activity is hypothesized to mitigate technical batch effects, its integration performance has never been quantitatively assessed against modern data integration benchmarks. We present flashscenic, an end-to-end GPU-accelerated framework that resolves these bottlenecks and expands the practical scope of single-cell regulatory genomics.

**Results:** By reformulating module construction, cisTarget motif pruning, and AUCell scoring into batched GPU tensor operations, flashscenic achieves a 12-to 27-fold downstream speedup over pySCENIC, while maintaining strict numerical equivalence (*r*_Pearson_ = 0.9996). Leveraging this throughput, we conducted the first systematic benchmark of nine consensus aggregation strategies across multiple independent tissue cohorts, demonstrating that discrete post-pruning recurrence voting (≥ 50% recurrence, ≥ 5 targets per regulon) maximizes biological fidelity. Finally, data integration benchmarking across a 116,043-cell acute myeloid leukemia (AML) atlas quantitatively demonstrates that uncorrected TF activity provides an intrinsic, unsupervised batch-integration benefit over raw PCA. Combining it with a standard batch integration method matches or outperforms canonical expression-based integration while accurately recovering lineage-defining and oncogenic regulators (CEBPA, HOXA9).

**Availability:** `pip install flashscenic`. Source code and tutorials: https://github.com/haozhu233/flashscenic.

## 1. Introduction

The regulation of gene expression programs underlies cellular development, differentiation, and tissue homeostasis (Garcia-Alonso et al. 2019). Central to this regulation are transcription factors (TFs), which link intracellular signaling pathways to gene-specific transcriptional control. Through these interactions, TFs establish gene regulatory networks (GRNs) that coordinate gene expression programs and ultimately define cellular identity and function. Disruption of these regulatory programs is a fundamental driver of disease, contributing to aberrant phenotypes in cancer (Gonzalez-Perez 2016, Osmanbeyoglu et al. 2017) degenerative disorders (Berson et al. 2018, Jin et al. 2019) and age-associated physiological decline (Levy et al. 2020, Yang et al. 2023). Consequently, reconstructing GRNs provides both mechanistic insight into disease processes and a framework for identifying candidate therapeutic targets.

The advent of single-cell RNA sequencing (scRNA-seq) has driven widespread adoption of GRN-based analysis frameworks, most prominently SCENIC (Aibar et al. 2017, Van de Sande et al. 2020). By projecting high-dimensional expression profiles into biologically interpretable transcription factor (TF) regulon activities, SCENIC has become a foundational tool for analyzing regulatory patterns across diverse biomedical contexts. However, while standalone GRN inference algorithms have rapidly advanced, spanning mutual information (Margolin et al. 2006, Chan et al. 2017, Qiu et al. 2020, Kim et al. 2021), decision tree models (Moerman et al. 2019), variational autoencoders (Shu et al. 2021, Zhu and Slonim 2025), and denoising diffusion models (Zhu and Slonim 2024), the overarching SCENIC workflow remains computationally constrained by sequential, CPU-bound cisTarget motif enrichment and per-cell AUCell scoring. These nested iteration bottlenecks frequently force studies to restrict analyses to narrow subsets of highly variable genes (HVGs) and render the multi-run ensembling needed to resolve stochastic network variability computationally prohibitive on large cohorts.

To address this limitation, we developed flashscenic, a GPU-accelerated framework that broadens the practical scope of motif-validated GRN analysis for large scale single-cell transcriptomics. In this study, our contributions are threefold:

- Vectorized GPU Architecture and Algorithmic Equivalence: flashscenic replaces key bottlenecks of the original pipeline with GPU native implementations, incorporating RegDiffusion for network inference together with fully vectorized PyTorch implementations of cisTarget pruning and AUCell scoring. Benchmarking shows it achieves a 12–27-fold downstream speedup over pySCENIC on a single GPU with verified numerical equivalence (*r*_Pearson_ = 0.9996). During validation, we also identified and corrected a systematic leading edge computation error in pySCENIC v0.12.1 that caused index misalignment in 98.9% of enriched motifs, producing more accurate regulon gene sets.
- Systematic Benchmark of Multi-Run Ensembling: Leveraging this computational throughput, we perform the first large-scale systematic evaluation of multi-run aggregation across three independent single-cell atlases, demonstrating that discrete post-pruning recurrence voting (≥ 50% recurrence, ≥ 5 genes per regulon) effectively resolves inherent GRN stochasticity and maximizes biological fidelity.
- Quantitative Assessment of Regulon Activity for Batch Integration: Using the standardized scIB benchmarking framework on a 116,043-cell AML bone marrow atlas, we provide the first quantitative evaluation showing that regulon-activity embeddings effectively reduce batch variation as an unsupervised substrate, and demonstrate that layering canonical correction (flashscenic + Harmony (Korsunsky et al. 2019)) matches or outperforms standard expression-based integration while recovering established leukemic regulatory programs.

## 2. Methods

### 2.1. Pipeline overview

At a high level, flashscenic follows the SCENIC pipeline (Aibar et al. 2017, Van de Sande et al. 2020). From a gene expression input matrix **X** ∈ ℝ^*n*×*p*^ (*n* cells, *p* genes), it produces a regulon activity matrix **A** ∈ ℝ^*n*×*r*^ (*r* regulons) in five stages: (1) GRN inference and TF filtering, (2) module construction (3) cisTarget motif-based pruning, (4) AUCell regulon activity scoring, and (5) regulon specificity scoring. Our key design principle is to reformulate each stage as batched tensor operations on the GPU, replacing the sequential per-module and per-cell loops used in pySCENIC (Van de Sande et al. 2020) with parallel tensor operations. (Figure 1). We use the term *module* to refer to a candidate TF–target gene set before motif validation, and *regulon* for the validated set after cisTarget pruning.

**Figure 1.**
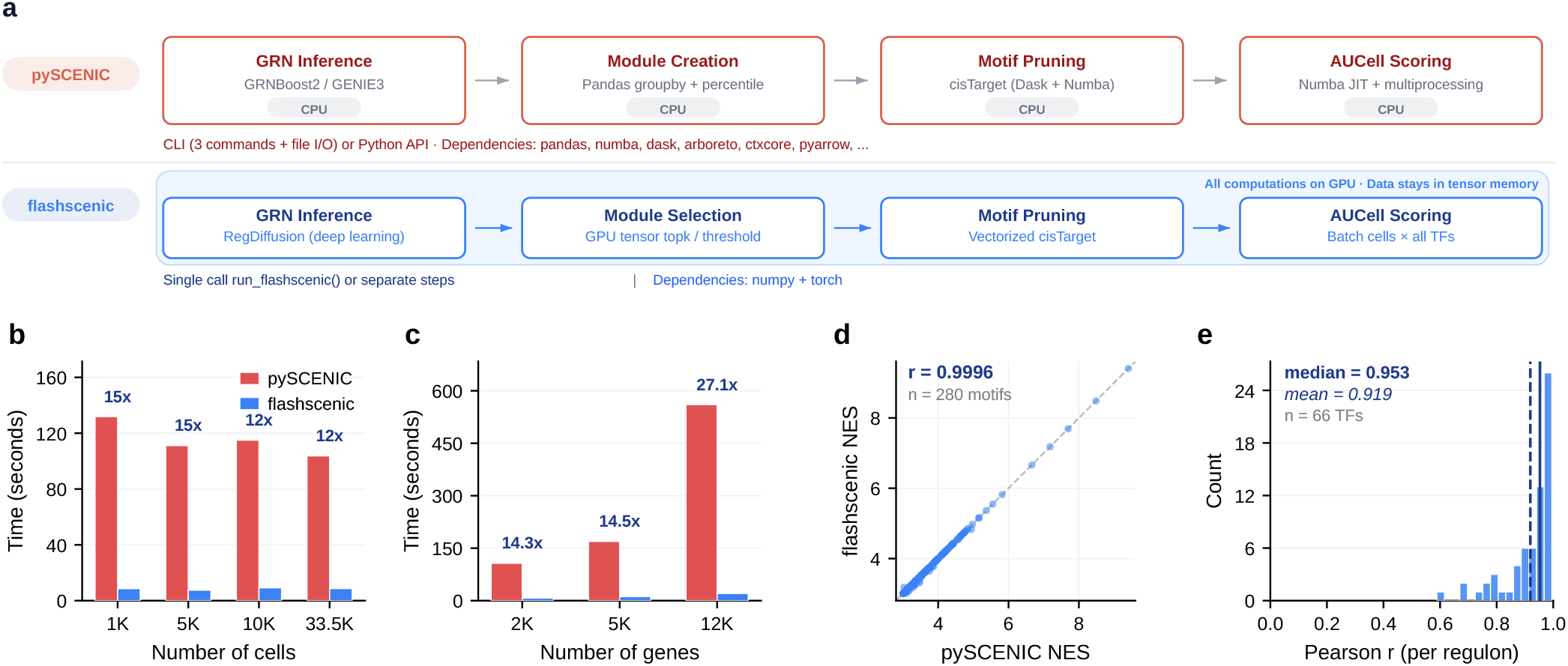
Algorithmic architecture, computational scalability, and numerical equivalence of flashscenic. **(a)** End-to-end pipeline comparison. PySCENIC operates on CPU with support of multi-processing (via Dask and Numba) while flashscenic replaces each stage with a GPU-native operation, keeping all data in tensor memory via a single function call. **(b)** Runtime Scalability as a function of number of cells, using 2,000 genes. **(c)** Runtime Scalability as a function of number of genes, using 10,000 cells. **(d)** Numeric equivalence on per-motif normalized enrichment scores (NES). **(e)** AUCell scores are highly correlated between pySCENIC and flashscenic.

#### GRN inference and TF filtering

We use RegDiffusion (Zhu and Slonim 2024) for GRN inference. Briefly, RegDiffusion trains a denoising structural equation model (SEM) on the expression matrix, where a learnable adjacency matrix **W** ∈ ℝ^*p*×*p*^ parameterizes gene–gene regulatory relationships. After training for *T* = 1,000 steps on GPU, we extract the learned adjacency matrix, where rows correspond to known transcription factors, yielding a TF-by-gene matrix **W**_TF_ ∈ ℝ^*m*×*p*^ (*m* TFs present in the data). Edges with weight below a sparsity threshold (*τ* = 1.0) are set to zero. The resulting sparse matrix serves as input for module construction.

#### Multi-strategy module construction

The SCENIC workflow constructs multiple candidate module types from the same adjacency matrix to increase sensitivity for regulon discovery (Aibar et al. 2017, Van de Sande et al. 2020). In pySCENIC, each module is represented as a GeneSignature object and constructed individually via list operations. flashscenic instead represents the full set of modules as a dense TF-by-gene tensor, allowing all module types and all TFs to be constructed simultaneously as parallel matrix operations. Following pySCENIC, we consider the following module types: (1) top-*k* targets per TF (default *k* = 50), (2) top-*N* TF per target module (*N* ∈ {5, 10, 50}), (3) percentile threshold modules (default 75%). Modules with fewer than 20 non-zero targets are discarded. Targets whose edge weights exceed a global percentile threshold are retained.

#### cisTarget pruning

CisTarget pruning (Imrichová et al. 2015) validates candidate modules by testing whether target genes are enriched for cis-regulatory motifs of the cognate TF. We use the v10 cisTarget ranking databases (Van de Sande et al. 2020), which provide genome-wide rankings of genes for each of ∼30,000 clustered motifs based on regulatory region scores. Two databases capturing different regulatory windows are used: a proximal database (500 bp upstream, 100 bp downstream of TSS) and a distal database (10 kbp flanking). Each database is represented as an integer-valued matrix **R** ∈ Z^*q*×*p*^ (*q* motifs, *p* genes), where *R*_*ij*_ is the rank of gene *j* for motif *i*. flashscenic loads the full database into GPU memory once and computes recovery curves for all *q* motifs simultaneously via vectorized scatter operations (up to a rank threshold *d* = 5, 000), rather than iterating per motif as in pySCENIC.

The area under each recovery curve (AUC) up to a cutoff rank *c* = ⌊0.05 · *p*⌋ − 1 is converted to a normalized enrichment score (NES) by z-scoring across all motifs, measuring the overrepresentation of a transcription factor’s target gene set within the top-ranked genes for a given motif. Specifically:

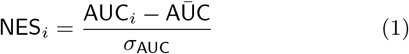

where AŪ C and *σ*_AUC_ are the mean and population standard deviation. Motifs with NES ≥ 3.0 are considered significantly enriched.

For each enriched motif, leading-edge genes are identified via a single broadcasted comparison against a background curve (defined as mean + 2· standard deviation across all motifs’ recovery curves), then linked to TFs via a motif annotation table (FDR *<* 0.001). When multiple databases are used, regulons are merged by union of leading-edge gene sets per TF.

#### AUCell scoring

AUCell (Aibar et al. 2017) quantifies regulon activity per cell by measuring whether a regulon’s target genes are enriched among the cell’s most highly expressed genes. In pySCENIC, AUCell iterates over regulons sequentially and, within each regulon, computes per-cell scores one at a time via a JIT-compiled function. flashscenic eliminates both loops by scoring all regulons and all cells in a batch simultaneously on the GPU. Per-cell gene rankings and target gene extraction are performed as batch matrix operations, and the AUC for each cell–regulon pair is computed through vectorized sorting and cumulative summation, with a rank cutoff at *c* = ⌊0.05 · *p*⌋ − 1 genes and per-regulon normalization by (*c* + 1) · _*j*_ *w*_*j*_, where *w*_*j*_ are target gene weights (binary in the standard pipeline). Cells are processed in configurable batches for GPU memory efficiency. The resulting activity matrix **A** ∈ ℝ^*n*×*r*^ provides a cell embedding that captures transcriptional regulatory state.

#### Regulon specificity scoring

To identify regulons with cell-type-specific activity, flashscenic computes Regulon Specificity Scores (RSS) (Suo et al. 2018) based on the Jensen–Shannon divergence (JSD). For each regulon *j* and cell type *t*, the AUCell score vector **a**_*j*_ ∈ ℝ^*n*^ is normalized to a probability distribution **â**_*j*_, and a binary indicator vector **1**_*t*_ (1 for cells of type *t*, 0 otherwise) is similarly normalized. The RSS is defined as 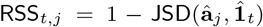, where values close to 1 indicate that the regulon’s activity is concentrated in that cell type. This provides a ranked list of regulons per cell type for downstream biological interpretation.

### 2.2. Benchmarking and equivalence

Datasets, filtering methods, and pre-processing methods are described in the Supplementary Methods and Supplementary Table ST1. flashscenic was run on a single NVIDIA A100 80GB GPU; pySCENIC v0.12.1 was run with 24 CPU cores.

Runtime benchmarking was performed on the Immune PBMC dataset. Cell-scaling experiments varied cell count (1,000–33,506) at 2,000 Highly Variable Genes (HVGs); gene-scaling experiments varied gene count (2,000–12,282) at 10,000 cells. To ensure fair comparison, we excluded the GRN inference time from the comparison. Instead, flashscenic and pySCENIC were both given the same RegDiffusion-inferred adjacency matrix and identical cisTarget parameters. Equivalence to pySCENIC was assessed via NES correlation across 280 commonly enriched (TF, motif, database) trios per-cell AUCell score correlation across 66 common TFs, and Jaccard overlap of regulon gene sets. During this analysis, we identified and corrected a systematic leading-edge bug in pySCENIC v0.12.1 (Supplementary Section S1.1); all comparisons use the patched version. Full benchmarking details are provided in Supplementary Section S1.2.

### 2.3. Multi-run aggregation strategy

Due to non-deterministic optimization and randomized mini-batch sampling, individual GRN inference runs produce variations in predicted edge weights that would propagate into downstream module selection and regulon scoring. To systematically quantify the impact of this run-to-run variation and establish an evidence-based aggregation method, we formalized and compared the following ensembling paradigms.

#### Pre-pruning continuous weight averaging

Continuous adjacency matrices from *n* independent GRN inference runs are merged into a single mean adjacency matrix, 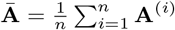. Unstable interactions can optionally be filtered out by applying a threshold on the coefficient of variance (CV = *σ*_**A**_*/*|**Ā** |, CV *<* 0.5*/*0.25*/*0.1). Once this consensus matrix is formed, the downstream module construction, motif pruning, and AUCell scoring steps are executed exactly once.

#### Post-pruning discrete frequency voting

An alternative is to run the entire five-stage pipeline independently across each of the *n* runs to produce *n* fully pruned regulon outputs. A final consensus is found by frequency voting on each TF-target pair, retaining only those that exceed a recurrence cutoff *f* (*f* ≥ 50%*/*75%*/*80%), with an optional minimum regulon size requirement.

We evaluated 9 consensus configurations (5 pre-pruning; 4 post-pruning) across *n* = 30 independent runs on three single-cell cohorts: ALS spinal cord, immune PBMC, and lung. Convergence was identified via the Jaccard similarity of TF-target pairs across run depths (*k* = 2, …, *n*). Activity stability was assessed by mean Spearman correlation of per-cell AUCell scores between regulons matched by TF name across consecutive *k* values. Biological accuracy was validated against CollecTRI (Müller-Dott et al. 2023), a manually curated resource that consolidates signed (activating/repressing) TF– target interactions from 12 databases into a single reference regulon set (52,281 interactions, 1,180 TFs, 8,457 targets). For each transcription factor, we computed precision recall, and F1 score, and we assessed enrichment with a one-sided Fisher’s exact test on the TF-target overlap. For each method, we carried out three independent runs of the full protocol with different seeds and reported the mean and standard deviation of recall and precision. Further details are provided in Supplementary Methods Sections S1.3 and S1.4.

### 2.4. AUCell for Batch integration

To quantitatively assess the potential of regulon activity as an unsupervised method for batch integration, we benchmarked flashscenic against conventional expression-based workflows using the Single-Cell Integration Benchmark (scIB) framework (Luecken et al. 2022). We first conducted a direct comparison between flashscenic and Raw PCA, since both are unsupervised methods with no batch information. We further compared these to two conventional single-cell data integration methods: Harmony (Korsunsky et al. 2019) and scVI (Lopez et al. 2018), where batch annotation is used to remove batch effects. Finally, we investigated applying Harmony in-place to the z-scored AUCell matrix (“flashscenic + Harmony”).

Integration performance was quantitatively assessed across 10 established scIB metrics. Five of these measure batch correction capacity: integration Local Inverse Simpson’s Index (iLISI), batch Average Silhouette Width (ASW batch), k-nearest-neighbor batch-effect test (kBET), Graph connectivity, and Principal Component Regression (PCR). The other five metrics measure biological conservation capacity: cell-type LISI (cLISI), Cell-type Average Silhouette Width (ASW cell type), Normalized Mutual Information (NMI), Adjusted Rand Index (ARI), and isolated label F1 score. Following the setup in scIB, the Total score was computed as the weighted average: 0.4 × batch score + 0.6 × bio conservation score. Hyperparameters, seed replicates, and distance matrix calculations are described in Supplementary Methods Section S1.5.

### 2.5. Datasets

The experiments above were conducted using four single-cell/single-nucleus RNA-seq cohorts spanning distinct cellular identities and scales. Immune PBMC (33,506 cells). The scIB immune reference atlas (Luecken et al. 2022) was used for computational scalability and numerical equivalence benchmarking against pySCENIC (Section 2.2). ALS spinal cord (62,711 nuclei), Immune PBMC, and Lung (32,281 cells). The ALS cohort (Takeuchi et al. 2025), together with the scIB immune and lung reference atlases (Luecken et al. 2022), was used to characterize multi-run stochasticity and benchmark consensus aggregation strategies (Section 2.3). AML bone marrow (116,043 cells; 4 studies; 36 donors). The multi-center AML cohort was used as the primary benchmark for regulon-activity-based batch integration, with the Immune PBMC and Lung atlases providing cross-tissue validation (Section 2.4; Supplementary Figs.S4–S5).

## 3. Results and Discussion

### 3.1. flashscenic improves scalability

To assess scalability, we benchmarked the downstream pipeline (module selection, cisTarget pruning, and AUCell scoring) against pySCENIC across varying cell numbers and gene set sizes on the Immune PBMC cohort (Figure 1b,c). Across 1,000 to 33,506 cells (at 2,000 HVGs), flashscenic executed downstream analysis in 7–9 seconds regardless of cell count, delivering a consistent 12-to 16-fold speedup over pySCENIC (104–132 s; Figure 1b). In particular, batched GPU tensor execution reduced AUCell scoring on 33,506 cells from 18.0 s to 0.14 s (*>*100× acceleration), outperforming specialized alternatives like UCell (Andreatta and Carmona 2021). When scaling gene set size from 2,000 to the full 12,282 genes (at 10,000 cells), flashscenic completed the pipeline in 8, 12, and 21 seconds, respectively, compared to 114, 170, and 561 seconds for pySCENIC, widening the speedup advantage from 14.3-fold to 27.1-fold (Figure 1c).

Beyond speed, scaling to the full transcriptome substantially expanded the number of inferred regulons: 66 regulons were identified from 2,000 HVGs, rising to 114 at 5,000 genes and 236 at the full 12,282-gene transcriptome. This near-quadrupling of recovered regulons suggests that standard HVG filtering, while computationally expedient, discards a substantial fraction of the regulatory signal detectable by motif enrichment analysis. TFs with tissue-specific or condition-specific expression patterns may be systematically under-represented among HVGs, which are selected on the basis of expression variance rather than regulatory relevance. The runtime improvements afforded by flashscenic therefore translate directly into expanded biological scope: analyses that were previously limited to a few thousand genes by computational cost can now routinely be applied to full transcriptomes.

### 3.2. Fidelity to SCENIC results

To confirm that GPU acceleration preserves numerical fidelity, we evaluated flashscenic against pySCENIC (using the patched version resolving the leading-edge indexing bug; Supplementary Section S1.1) on a shared RegDiffusion network across 10,000 Immune cells. Across 280 significantly enriched (TF, motif) pairs (*NES* ≥ 3.0), NES showed near-perfect agreement (Pearson *r* = 0.9996; Figure 1d), with 92% of pairs (258/280) displaying absolute NES differences below 10^−6^ due solely to minor floating-point differences in vectorized calculations. Downstream AUCell regulon activity scores were similarly concordant across 66 shared TFs, yielding a median per-TF Pearson correlation of 0.953 (mean *r* = 0.919; Figure 1e). Overall, 61% of regulons (40/66) produced gene-for-gene identical target sets (Jaccard index = 1.0; mean Jaccard = 0.945). The small fraction of TFs with lower agreement (e.g., STAT1, *r* = 0.589; STAT4, *r* = 0.742) tend to have small or borderline regulons where minor differences in how percentile-threshold and per-target module types are constructed have a larger impact. Overall, this level of agreement confirms that the core recovery curve and AUC computation is implemented identically in both systems at GPU and CPU precision.

### 3.3. Ensembling for GRN stability

#### Addressing stochasticity in GRN inference

The stochastic nature of GRN inference is a well-documented challenge in single-cell transcriptomics. As noted by the original pySCENIC paper (Van de Sande et al. 2020), decision tree based GRN inference methods, such as GENIE3 and GRNBoost2, rely on repeated random subsamplings, resulting in small run-to-run variations in network edge weights. These variations pass down through the SCENIC pipeline and shift the target regulon sets. The recommended solution for pySCENIC is to run the full pipeline multiple times and filter regulons by frequency. However, executing the full SCENIC pipeline repeatedly has proven computationally prohibitive for many users. With the runtime improvements of RegDiffusion and flashscenic’s GPU acceleration, this consensus approach is now highly practical. However, a systematic quantitative evaluation of the optimal aggregation strategy has not yet been explored in the literature.

#### Frequency-based voting yields the most accurate consensus regulons

To determine the most robust approach for handling this stochasticity, we systematically compared nine aggregation strategies across three independent single-cell datasets (Figure 2a). We validated the results against CollecTRI, a manually curated TF-target database, and we found that aggregating regulons after pruning via frequency voting consistently yields a higher number of validated TF-target gene, outperforming strategies that averaged continuous edge weights before pruning. Specifically, keeping TF-target pairs that appear in at least 50% of runs, together with a minimum size filter of 5 genes per regulon (Freq ≥ 50% min5g), was the best strategy, achieving the highest recall and F1 scores across all three datasets (ALS, Immune, and Lung). Compared to the stricter, pySCENIC-style 80% threshold, this lower threshold sits at a Pareto-favorable high-recall end of the precision/recall trade-off and captures more biologically valid TFs without severely compromising precision. Independent Fisher’s exact validation corroborates Freq ≥ 50% min5g as the best-performing strategy. As a complementary, per-TF check that does not depend on the TF-score ranking or the precision/recall comparison above, this approach recovered the highest fraction of significantly enriched TFs across most evaluated conditions (Supplementary Figure S2).

**Figure 2.**
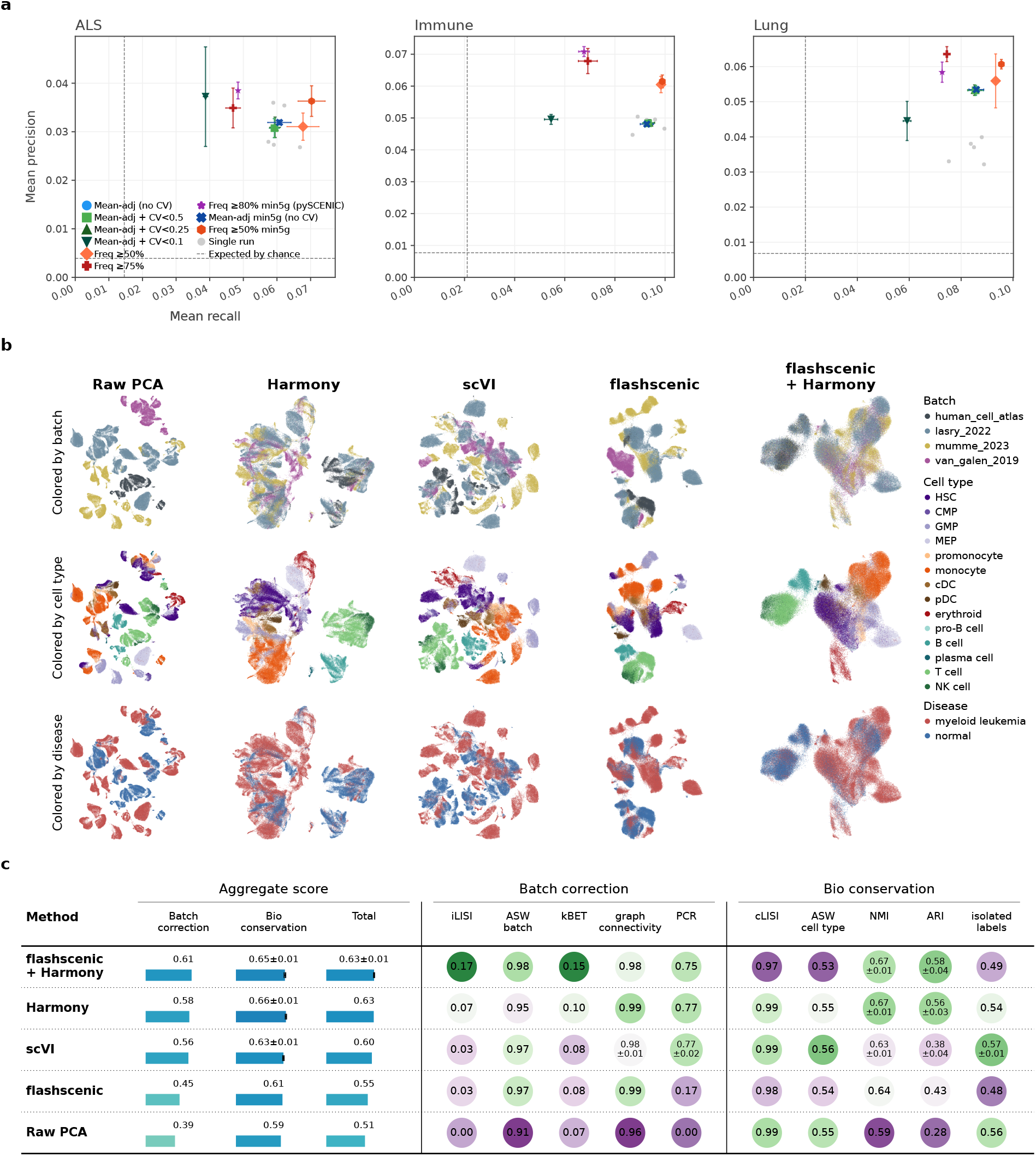
Multi-run consensus benchmarking and regulon-activity-based batch integration. **(a)** Precision and recall of nine consensus aggregation strategies (*k* = 30 runs) evaluated against the curated CollecTRI reference database across ALS, immune PBMC, and lung cohorts. Error bars represent standard deviations across three independent seed replicates of the full pipeline. **(b)** UMAP projections of 116,043 single-cell transcriptomes from the AML bone marrow atlas across five embedding representations (columns), colored by batch source, cell type, and disease status (rows). **(c)** Quantitative integration performance across 10 scIB metrics. Table cells are color-coded by column-wise rank (green: best, purple: worst), with aggregate and individual scores reported as mean *±* SD across five replicates for stochastic methods.

#### Post-pruning aggregation converges faster

Beyond biological accuracy, we also assessed how quickly each strategy converged to a stable consensus over 30 independent runs (Supplementary Fig. S1a,b). Using both Jaccard similarity on regulons and Spearman correlation of AUCell scores, post-pruning frequency voting converged significantly faster than mean-adjacency averaging. Frequency-based methods essentially converge within ∼15 runs, whereas mean-adjacency strategies often require 25-30. Further, freqency-based methods reached the highest AUCell correlation of all tested methods.

#### Size filtering reduces noise

While frequency-voting converges quickly, unfiltered baselines generate many small, under-supported regulons (Supplementary Fig. S1d). Applying the minimum 5-target-gene filter (the dashed line in Supplementary Fig. S1d) successfully eliminates these. At the same time, as shown in Supplementary Fig. S1c, the Freq ≥ 50% min5g strategy stabilizes the number of regulons to around 125 on this data set. Compared to the Freq ≥ 75% strategy, there are a similar number of regulons, but they tend to be larger and more evidence-supported, explaining the high recall and top-tier precision of the Freq ≥ 50% min5g strategy.

#### Strong edges are inherently stable, making CV filtering ineffective

Compared to frequency-based filters, mean-adjacency aggregation methods converge more slowly and underperform. Such methods were suggested for both DeepSEM (Shu et al. 2021) and RegDiffusion (Zhu and Slonim 2024). Their weak performance here stems from how individual edge weights vary across RegDiffusion runs. An intuitive way to address this variability might be to filter the averaged prediced edge weights by their coefficient of variation (CV). However, our experiments showed this approach to be ineffective (Supplementary Fig. S3). For the strongest edges, the absolute run-to-run variances eventually stabilize into a bounded additive-noise regime. Because every edge retained after the pipeline’s standard hard sparsity threshold already falls in this stable region, further filtering the averaged adjacency matrix based on CV loses its discriminatory power: mild CV thresholds do nothing, while aggressive CV thresholds discard real signal.

### 3.4. Batch integration; AML validation

#### Regulon-activity for unsupervised batch integration

The original SCENIC workflow (Aibar et al. 2017) demonstrated that scoring network activity is robust against technical variation and may be used to automatically overcome batch effects. However, this property has historically been observed qualitatively via dimension reduction and visualization rather than measured against modern integration metrics. To our knowledge, the present study provides the first quantitative evaluation of regulon activity for batch integration under modern benchmarking standards. Using the scIB evaluation framework across the 116,043-cell AML cohort and reference atlases, we benchmarked uncorrected flashscenic AUCell embeddings directly against raw expression PCA without providing batch labels (Figure 2b,c, Supplementary Figs. S4–S5). Across all evaluated cohorts, flashscenic activity improved batch correction scores over raw PCA while maintaining superior biological conservation. Because regulon activity scores are computed from motif-validated, co-regulated gene sets, this representation effectively collapses platform- and depth-specific technical noise while preserving genuine cell-type structure.

#### Continuous correction on regulon embeddings matches or exceeds state-of-the-art workflows

We next evaluated whether flashscenic embeddings could serve as an effective substrate for canonical batch correction algorithms. Because AUCell scoring produces a continuous latent matrix, Harmony was applied directly to the z-scored regulon activity space (flashscenic + Harmony). This strategy matched or exceeded expression-based Harmony across all three datasets in aggregate (Total) integration score, significantly so in lung (Figure 2b,c, Supplementary Figs. S4–S5, Supplementary Table ST3). These results demonstrate that motif-constrained TF activities provide a reliable, noise-resilient alternative input for downstream data integration.

#### flashscenic regulons recapitulate established leukemic circuitry and lineage programs

Further supporting the biological validity and applicability of flashscenic’s large-scale GRNs, downstream analyses recovered both established AML regulatory programs and coherent lineage-specific patterns. Differential TF activity (per cell type; Supplementary Fig. S6) reproduced established oncogenic regulators: **CEBPA** elevated in hematopoietic stem cells (HSC; Δ = +0.026, *q* = 0.003) and **HOXA9** elevated in monocytes (Δ = +0.007, *q* = 0.003), both centrally implicated in myeloid leukemogenesis (Collins et al. 2014, Sun et al. 2018). Notably, this elevation of HOXA9 activity coincided with reduced activity of the paralogous regulons **HOXA5** and **HOXA6** within the MEP compartment (Δ = −0.027, *q* = 0.034 and Δ = −0.057, *q* = 0.058, respectively), further implicating the HOXA regulatory axis, frequently dysregulated in AML (Alharbi et al. 2013).Independently, regulon specificity scores (RSS; Supplementary Fig. S7) confirmed that flashscenic regulons capture lineage-specific regulatory programs, recovering established regulators including **ERG** in HSCs (Knudsen et al. 2015), **NFE2** in MEPs (Shivdasani et al. 1995), **LEF1** and **TBX21** in T cells (Okamura et al. 1998).

### 3.5. Discussion

The results presented here establish flashscenic as a scalable and biologically interpretable tool for regulatory network analysis of large single-cell datasets. flashscenic addresses two practical bottlenecks in regulatory network inference at cohort scale: computational cost and the stochasticity inherent to any GRN-inference method built on randomized training. Its GPU-accelerated pruning and AUCell scoring make full-transcriptome regulon inference practical rather than the HVG-restricted analyses forced by pySCENIC’s runtime, nearly quadrupling the number of detectable regulons in our benchmarks. The consensus-aggregation analysis further shows that, once GRN inference is affordable to repeat, applying a less restrictive frequency threshold to pruned regulons, rather than averaging continuous edge weights before pruning, converges faster and recovers more regulators. This is consistent with our finding that edges retained by the pipeline’s default sparsity threshold exhibit relatively stable weights across runs, suggesting that post-pruning edge recurrence captures the remaining stochasticity more effectively than further averaging of continuous edge weights. This result held across three independent tissues, supporting Freq ≥ 50% min5g and *k* = 15 as a practical default for multi-run consensus GRN construction.

Our findings also clarify the utility of TF activity as an unsupervised method for data integration. Because regulon activity scores are derived from motif-validated, multi-gene consensus targets, they inherently dampen technical dropout and sequencing-depth disparities while preserving cellular identity. When combined with continuous correction algorithms like Harmony, flashscenic-derived activity matrices yield integrated embeddings that match or exceed conventional gene-expression workflows, all while providing directly interpretable regulatory states rather than abstract latent dimensions. However, because AUCell generates continuous enrichment scores rather than raw count distributions, it naturally pairs with continuous embedding correctors rather than count-based generative architectures like scVI, highlighting the need to match integration methods to data representation geometries.

Beyond flashscenic itself, this analysis surfaced a previously unreported bug in pySCENIC’s leading-edge computation (Supplementary Section S1.1): a DataFrame sort used to resolve motif-annotation ties silently desynchronizes the sorted annotation table from the arrays used to compute leading-edge target genes, corrupting the reported regulon gene sets for 98.9% of enriched motifs while leaving NES and AUC scores unaffected. We have reported this issue upstream; because it is present in all public pySCENIC releases through v0.12.1, prior studies that relied on pySCENIC’s leading-edge output for downstream regulon interpretation may warrant reanalysis.

## 4. Conclusions

We presented flashscenic, a GPU-accelerated implementation of the SCENIC pipeline that resolves the scalability, reproducibility, and stability bottlenecks of single-cell gene regulatory network analysis. By replacing sequential CPU loops with fully vectorized tensor operations, flashscenic achieves 12–27-fold downstream speedups with numerical equivalence, lowering the barrier to large-scale GRN inference. Beyond the runtime contribution, we established an evidence-based multi-run ensembling strategy (Freq ≥ 50% min5g) that effectively stabilizes stochastic networks, and demonstrated that regulon activity serves as a robust unsupervised batch integration method. Overall, flashscenic broadens the practical applicability of the SCENIC approach to transcriptome-wide cohort-scale scRNAseq studies where computational efficiency and downstream interpretability are both required.

## Supporting information

supplement doc

## 5. Author contributions

H.Z., D.K.S., and X.J. drafted the ideas; H.Z. and G.C.A. implemented the code; G.C.A., H.Z. and X.J. conducted the experiments; G.C.A., H.Z., and D.K.S. analyzed the results. H.Z., G.C.A., and D.K.S. wrote the initial draft. All authors reviewed and edited the manuscript.

## 6. Code availability

flashscenic can be installed via pip install flashscenic. Source code, experiments and tutorials are available at: https://github.com/haozhu233/flashscenic.

## Notes

https://github.com/haozhu233/flashscenic

### Competing Interest Statement

The authors have declared no competing interest.

