## supplement doc for "flashscenic: Accelerated GRN Analysis with RegDiffusion for Large-Scale Single-Cell Data"

Supplement for flashscenic: Accelerated gene regulatory network inference with RegDiffusion for large-scale single-cell data

### 1 Supplemental Methods

#### 1.1 Identification and correction of the pySCENIC leading-edge bug

During equivalence analysis, poor regulon gene set agreement despite high NES concordance prompted investigation of the leading-edge computation in pySCENIC v0.12.1. The root cause was identified in `pyscenic/transform.py`, function `module2features_auc1st_impl`: following identification of enriched motifs and motif annotation, the DataFrame is sorted by `MotifSimilarityQvalue` and `OrthologousIdentity` to resolve ties, changing row order. A subsequently computed boolean annotation mask is then applied to recovery curve (`rccs`) and gene ranking (`rankings`) arrays that retain the original, pre-sort order, causing systematic index misalignment. The bug was corrected by inserting `annotated_features = annotated_features.reindex(enriched_features.index)` immediately after the sort-and-deduplication step to restore alignment before mask application. This patch was applied to both the local pySCENIC source and the installed package environment prior to all equivalence analyses and benchmarks reported here.

#### 1.2 Runtime benchmarking details

Runtime benchmarking and equivalence analyses were performed on the scIB human immune PBMC benchmark atlas, containing 33,506 cells from 10 batches spanning multiple technologies and donors, 16 cell types). The same raw dataset is underlying the batch-integration and multi-run robustness analyses (Section 1.3.4) but preprocessed independently here. Cell-scaling experiments used successive subsamples of 1,000, 5,000, 10,000, and the full 33,506 cells; all remaining experiments (gene-scaling, NES equivalence, AUCell equivalence, and regulon gene set overlap) were run on a fixed subsample of 10,000 cells. Gene filtering and highly-variable gene (HVG) selection were performed in Scanpy under two regimes: an HVG regime, in which genes were selected with `sc.pp.highly_variable_genes` (`flavor='cell_ranger'`) at a target count of 2,000 or 5,000 genes, and a “full-transcriptome” regime, in which all genes detected in more than 10 cells were retained (resulting in 12,282 genes after filtering). In both regimes, expression matrices were library-size normalized to 10,000 counts per cell and log1p-transformed prior to GRN inference.

flashscenic computations were performed on a single NVIDIA A100-SXM4-80GB GPU (CUDA) with 24 CPU cores. pySCENIC v0.12.1 was run using the command-line interface (`pyscenic ctx` and `pyscenic aucell`) with 24 parallel CPU workers. Both tools used the same GRN adjacency matrix as input. The hg38 v10 cisTarget ranking databases (500 bp upstream/100 bp downstream and 10 kbp upstream/10 kbp downstream of the transcription start site) were used for motif pruning, with the corresponding motif annotation table (`motifs-v10nr_clust-nr.hgnc-m0.001-o0.0.tbl`).

Wall-clock time was measured using Python’s `time.time()` function, recording the elapsed time of each pipeline stage independently. For pySCENIC, timing was measured around the respective CLI subprocess calls. GRN inference time was removed from the comparison. Specifically, for both cell-scaling and gene-scaling comparisons, only downstream time (module selection + cisTarget + AUCell) was compared, since GRN inference in RegDiffusion is known to be faster than comparable inference steps with GENIE3 or GRNBoost2 (Zhu and Slonim, 2024).

For equivalence analysis, both tools used identical pruning parameters: `rank_threshold` = 5,000, `auc_threshold` = 0.05, `nes_threshold` = 3.0, with both cisTarget databases (500 bp and 10 kbp), on a shared RegDiffusion adjacency matrix (seed = 42 for cell selection) at 10,000 cells and 2,000 HVGs. NES correlation was computed across all commonly enriched (TF, motif, database) pairs; AUCell correlation was measured per TF across cells; Jaccard overlap was computed per TF between regulon gene sets.

#### 1.3 Data preprocessing

##### 1.3.1 Shared quality control and normalization

All four datasets described below (AML, ALS spinal cord, and the scIB immune and lung benchmark atlases; **Table S1**) were quality-controlled and normalized with the same thresholds. Raw counts were identified automatically from `adata.X`, `adata.raw`, or a counts layer, and Ensembl gene IDs (where present) were mapped to HGNC symbols. Mitochondrial content was computed from `MT-`-prefixed genes; cells were retained if they expressed $\geq 200$ genes and had $\leq 20\%$ mitochondrial counts. Genes were retained if detected in at least 1% of cells. Counts were library-size normalized to 10,000 and log1p-transformed; no highly-variable-gene subsetting was applied, so all QC-passing genes were carried forward into GRN inference and regulon scoring.

**Table S1.** Summary of data sets used. SC = single cell; SN = single nucleus. Data set sizes are after filtering for QC only, and do not capture further downsampling of data set sizes described in the main Methods text for some analyses. Analyses are referred to as Scalability (Main paper Results section 3.1), Fidelity (3.2), Stochasticity (3.3), and Harmonization (3.4).

| Name | Reference | Batches | Type | Cells/Nuclei | Genes | Analyses |
|---|---|---|---|---|---|---|
| AML | (Whittle et al., 2026) | 4 | SC | 116,043 | 10,740 | Harmonization |
| ALS | (Takeuchi et al., 2025) | 1 | SN | 62,711 | 17,994 | Stochasticity |
| Immune | (Luecken et al., 2022) | 10 | SC | 33,506 | 10,410 | Scalability<br>Stochasticity<br>Harmonization |
| Lung | (Luecken et al., 2022) | 16 | SC | 32,281 | 13,184 | Stochasticity<br>Harmonization |

##### 1.3.2 ALS spinal cord dataset

We applied flashscenic to 62,711 single-nucleus RNA-seq profiles from human ALS and healthy control spinal cord, yielding 17,994 genes after QC. This dataset is the primary substrate for the multi-run GRN-inference robustness analysis and for the RegDiffusion stochasticity characterization below (Section 1.4, Supplementary Fig. S3).

##### 1.3.3 AML dataset

We additionally applied flashscenic to a publicly available bone marrow scRNA-seq atlas spanning AML and healthy donors (Whittle et al., 2026), comprising four studies: two contributing both AML and normal donors (*van_galen_2019*, *lasry_2022*, 5 normal donors each), supplemented with one normal-only study (*human_cell_atlas*, 8 donors) and one AML-only study (*mumme_2023*, 8 donors) to reach a balanced 18 AML / 18 normal donors (36 total), via deterministic top-donor-count-by-cell-count selection within each study. `Study` was used as the batch-correction covariate. After quality control, this yielded 116,043 cells and 10,740 genes across 14 annotated bone marrow cell types.

##### 1.3.4 Immune PBMC and lung benchmark datasets

The scIB human immune PBMC (33,506 cells, 10,410 genes after QC, 10 batches, 16 cell types) and human lung (32,281 cells, 13,184 genes after QC, 16 batches, 17 cell types) atlases (Luecken et al., 2022) were used as additional validation datasets in both independent analyses: as generalization checks for the multi-run robustness analysis, Supplementary Figs. S2–S3), and as generalization checks for the AML batch-integration case study (Supplementary Figs. S4–S5).

#### 1.4 Characterisation of RegDiffusion stochasticity

To characterise the run-to-run stochasticity of RegDiffusion GRN inference and its relationship to flashscenic's edge-filtering step, RegDiffusion was run $k = 30$ times independently (identical hyperparameters as single-run GRN inference, $T = 1{,}000$ gradient steps each) on a fixed, previously preprocessed expression matrix, yielding adjacency matrices $\mathbf{A}^{(1)}, \ldots, \mathbf{A}^{(k)}$ over the same gene set. The per-edge consensus mean $\bar{\mathbf{A}} = \frac{1}{k} \sum_i \mathbf{A}^{(i)}$ and standard deviation $\sigma_{\mathbf{A}}$ across runs were computed, along with the per-edge coefficient of variation $\mathrm{CV} = \sigma_{\mathbf{A}} / |\bar{\mathbf{A}}|$ (set to 0 where $\bar{\mathbf{A}} = 0$). Only edges with $\bar{\mathbf{A}} > 0$ (i.e. edges with non-zero average support across runs) were retained for all three analyses below. This procedure was applied independently to three datasets: the ALS spinal cord cohort (Section 1.3.2) and the scIB immune and lung robustness datasets. As all three analyses below yielded qualitatively consistent results across every dataset examined, only the ALS run is reported in full, shown directly in Supplementary Fig. S3 and quoted in the main text, with consistency against the immune and lung datasets confirmed separately.

##### 1.4.1 Run-to-consensus deviation (Bland–Altman-style scatter)

For each of the $k = 30$ runs and each retained edge, the deviation from consensus, $\mathbf{A}_e^{(i)} - \bar{\mathbf{A}}_e$, was computed and plotted against the consensus edge weight $\bar{\mathbf{A}}_e$ (Supplementary Fig. S3a). To summarise the envelope of variability as a function of edge strength without assuming a parametric noise model, consensus weights were partitioned into 50 bins by empirical percentile (bin edges at the 0th, 2nd, ..., 100th percentiles of $\bar{\mathbf{A}}_e$; bins containing fewer than 5 edges were dropped), and the empirical 5th and 95th percentiles of the pooled per-run deviations within each bin were computed and rendered as a shaded band. Because plotting every (edge, run) pair is impractical, scatter points were subsampled without replacement (maximum $5 \times 10^5$ points, seed 0) prior to plotting; the percentile bands themselves were computed from the full, non-subsampled set of deviations.

##### 1.4.2 Edge-weight coefficient-of-variation distribution

For the same retained edges, the distribution of per-edge $\mathrm{CV}_e$ across the $k$ runs was summarised as an 80-bin histogram (Supplementary Fig. S3b). Three CV thresholds (0.5, 0.25, 0.1), corresponding to the CV-based consensus-filtering strategies compared in main-text Fig. 3, were overlaid as vertical reference lines, and the fraction of edges with $\mathrm{CV}_e$ at or above each threshold was computed directly from the per-edge CV values and annotated on the plot.

##### 1.4.3 Weight–CV equivalence and the sparsity threshold

To relate this run-to-run CV to flashscenic's actual edge-filtering criterion, edges were sorted by consensus weight $\bar{\mathbf{A}}_e$ and partitioned into 19 log-spaced bins spanning the observed weight range (bins with fewer than 5 edges were discarded); the median CV within each retained bin was computed as a function of the bin's median weight and plotted on log–log axes (Supplementary Fig. S3c). The CV implicitly enforced by flashscenic's hard sparsity threshold ($\tau = 1.5$, the default edge-weight cutoff below which edges are zeroed prior to module construction; Section 1.3.2) was obtained by linear interpolation between the full, unbinned per-edge weight and CV values at weight $= \tau$; this value is marked with a dashed vertical line and annotated on the panel ($\mathrm{CV} \approx 0.16$ for the ALS run shown; also reported in the main text). This interpolated value is markedly lower than any of the three CV thresholds swept for the mean-adjacency+CV consensus strategies in main-text Fig. 3: the pipeline's hard weight cutoff already enforces a substantially stricter effective CV than any threshold subsequently swept, which explains why sweeping those thresholds changes regulon counts little until the most aggressive setting is reached ($\mathrm{CV}<0.1$).

#### 1.5 Batch Integration and Differential TF Activity in AML

##### 1.5.1 Cell embedding and integration quality

We compared five cell representations for their ability to remove batch/study effects while preserving cell-type and biological structure: (1) raw PCA (50 components, uncorrected,*Raw PCA*); (2) Harmony (Korsunsky et al., 2019) applied to raw PCA (*Harmony*); (3) scVI (Lopez et al., 2018) applied to raw counts, with disease status included as a categorical covariate for AML, where disease labels are available (*scVI*); (4) the flashscenic regulon-activity AUCell matrix, uncorrected (*flashscenic*); and (5) Harmony applied directly to the z-scored flashscenic regulon-activity matrix (*flashscenic+Harmony*). Harmony was run with `max_iter_harmony` = 20, with `theta` and `lambda` left at harmonypy's library defaults (1 and 1, respectively); scVI was run with `n_latent` = 10, `n_layers` = 1, `early_stopping` = `False`, and `max_epochs` = `None`.

Integration quality was assessed with the standard scIB metric set: five batch-removal metrics (iLISI, $ASW_{batch}$, kBET, graph connectivity, principal component regression [PCR] batch-variance reduction relative to the raw-PCA baseline) and five biological-conservation metrics (cLISI, $ASW_{cell\ type}$, NMI and ARI of Leiden clustering against annotated cell type at four resolutions [0.1, 0.3, 0.5, 1.0], and the isolated-labels ASW score). iLISI, kBET, graph connectivity, and cLISI were computed on a $k = 90$ neighbor graph; NMI and ARI's Leiden clustering used a separate $k = 15$ neighbor graph, following standard practice for graph-based clustering (a wide neighborhood as used for the LISI-family metrics would over-smooth fine cell-type boundaries). Per-metric scores were combined into `batch_score` (mean of the five batch-removal metrics) and `bio_score` (mean of the five bio-conservation metrics), and into a single `Total` score as $0.4 \times \texttt{batch\_score} + 0.6 \times \texttt{bio\_score}$, following the scIB convention.

Three of the five representations involve stochastic fitting – Harmony's k-means-based clustering initialization and scVI's neural-network training – and are correspondingly sensitive to random seed. For *Harmony*, *flashscenic+Harmony*, and *scVI*, we therefore repeated the fit five times per dataset at five independent random seeds (42–46) and report the mean ± standard deviation of each metric across replicates. *Raw PCA* and *flashscenic* (the two uncorrected representations) involve no stochastic fitting step and are reported as single values. Differences in `Total` score between every pair of the three seed-replicated methods were tested with a two-sample Welch's $t$-test (unequal variance, 5 vs. 5 observations), and Benjamini–Hochberg corrected across the three pairwise comparisons within each dataset. To test generality beyond AML, the identical protocol was independently repeated on the same two public scIB benchmark atlases.

##### 1.5.2 Differential TF activity analysis

To identify TF regulons with disease-associated activity within each cell type, we fit, for every regulon and every cell type with $\geq 500$ cells, a zero-inflated Beta mixed model (glmmTMB (Brooks et al., 2017)) of per-cell AUCell activity as a function of disease status, with donor as a random intercept in both the conditional and zero-inflation components: $\texttt{AUC} \sim \texttt{disease} + \texttt{sex} + (1 \mid \texttt{donor})$, zero-inflation $\sim 1 + (1 \mid \texttt{donor})$. $p$-values for the disease fixed effect were Benjamini–Hochberg corrected within each cell type. We report the effect size as the mean AUCell activity difference between disease and control cells, $\Delta = \overline{\texttt{AUC}}_{\text{disease}} - \overline{\texttt{AUC}}_{\text{control}}$, and visualize the top 10 regulons per cell type by $|\Delta|$ (colored by direction) as a heatmap (Fig. S6).

##### 1.5.3 Cell-type-specific regulon analysis

Cell-type-specific regulons were identified using the RSS. The top five regulons per cell type by RSS were visualised as a heatmap S7.

### 2 Supplemental Results

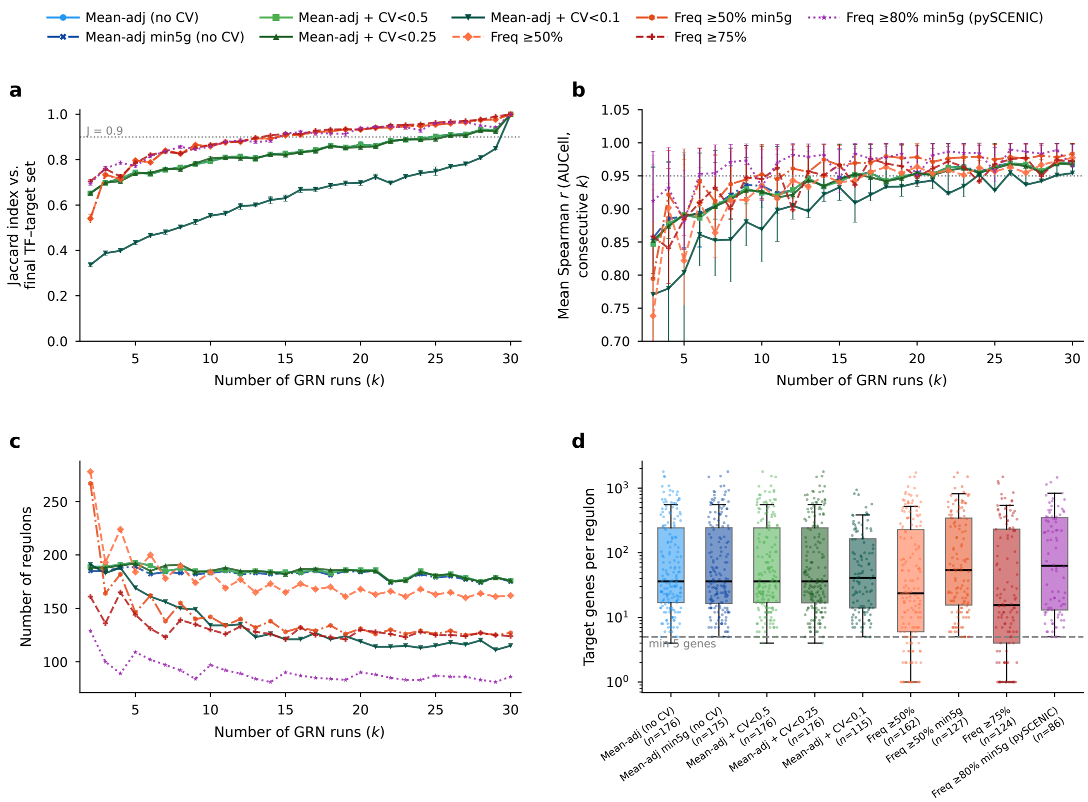


**Fig S1. Convergence of consensus aggregation strategies with increasing GRN-inference run depth, ALS spinal cord cohort.** All nine consensus aggregation strategies (main text Fig. 2a) evaluated over $k = 2$ to 30 independent RegDiffusion runs on the ALS spinal cord cohort (62,711 nuclei, 17,994 genes after QC). **(a)** Jaccard similarity between the consensus TF-target gene set at each $k$ and the final ($k = 30$) consensus set, as a function of $k$. **(b)** Mean Spearman correlation of per-TF AUCell activity scores between consensus regulon sets built from consecutive values of $k$. Frequency-based post-pruning strategies (orange/red/purple) converge to high similarity with their final regulon set within about 15 runs, whereas mean-adjacency averaging strategies (blue/green) require 25–30 runs to reach comparable similarity. **(c)** Number of regulons retained as a function of $k$. **(d)** Distribution of target genes per regulon at $k = 30$ for each strategy (points: individual regulons; box: median and interquartile range); the dashed line marks the 5-target-gene minimum-size filter. Unfiltered mean-adjacency baselines (blue/green) retain many small, weakly supported regulons, while frequency-based strategies with the min5g filter (orange/red/purple) are shifted toward larger, better-supported regulons. Freq ≥50% min5g (main text default) reaches high run-to-run similarity fastest among the strategies that are not overly aggressive, and stabilizes to approximately 125 regulons of predominantly larger size.

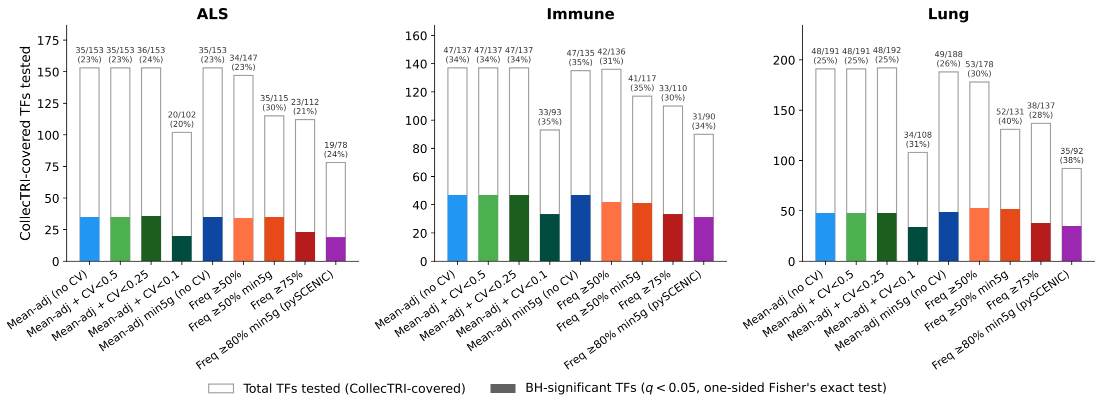


**Fig S2. CollecTRI Fisher's-exact validation of the nine consensus aggregation strategies, per dataset.** For each of the nine $k = 30$ consensus aggregation strategies and each of the three robustness datasets (ALS spinal cord, scIB immune, scIB lung), the open (outline-only) bar shows the total number of CollecTRI-covered transcription factors tested, and the filled bar (strategy's own colour, matching Fig. 3 in the main text) shows the number that are BH-significant ($q < 0.05$, one-sided Fisher's exact test for enrichment of the predicted target set against the CollecTRI reference, relative to the gene universe). The label above each bar pair gives the significant/tested count and its percentage. Freq ≥50% min5g attains the highest significant fraction of all nine strategies in ALS (35/115, 30.4%) and lung (52/131, 39.7%), and the second highest in immune (41/117, 35.0%), while also testing more TFs than the stricter Freq ≥80% min5g (pySCENIC-style) baseline in every dataset.

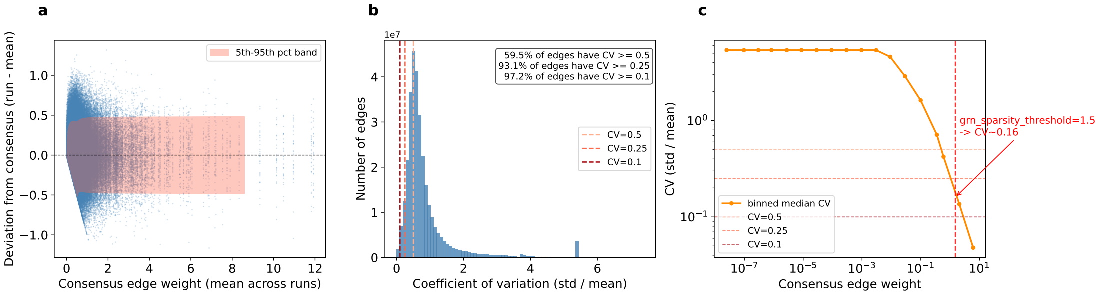


**Fig S3. Run-to-run edge-weight variability of RegDiffusion GRN inference, and its relationship to the pipeline's hard sparsity threshold.** All three panels are shown for the ALS spinal cord cohort (main text, *Consensus aggregation stabilizes GRN inference across runs*; 62,711 nuclei, 17,994 genes after QC, 30 RegDiffusion runs), the same dataset the corresponding main-text values (panels a–b) are quoted from, since the qualitative shape of all three panels was consistent across every dataset examined, including immune and lung, checked separately but not shown. **(a)** Bland–Altman-style scatter of each run's deviation from the 30-run consensus edge weight ($\mathbf{A}^{(i)} - \bar{\mathbf{A}}$) as a function of consensus edge weight; the shaded band is the empirical 5th–95th percentile envelope over 50 weight bins. **(b)** Distribution of per-edge coefficients of variation ($\mathrm{CV} = \sigma_{\mathbf{A}}/|\bar{\mathbf{A}}|$) across the 30 runs, with the fraction of edges exceeding each of the three CV thresholds used by the mean-adj+CV aggregation strategies (main text) annotated. **(c)** Median CV as a function of consensus edge weight (binned, log–log axes), showing that CV falls off approximately as the inverse of edge weight; the red dashed line marks the pipeline's hard edge-weight sparsity threshold ($\tau = 1.5$, used to binarize the adjacency matrix upstream of module construction) and the implied CV it already enforces, explaining why the CV-masking strategies swept in the main text's Fig. 3 change little until the most aggressive threshold (CV<0.1).

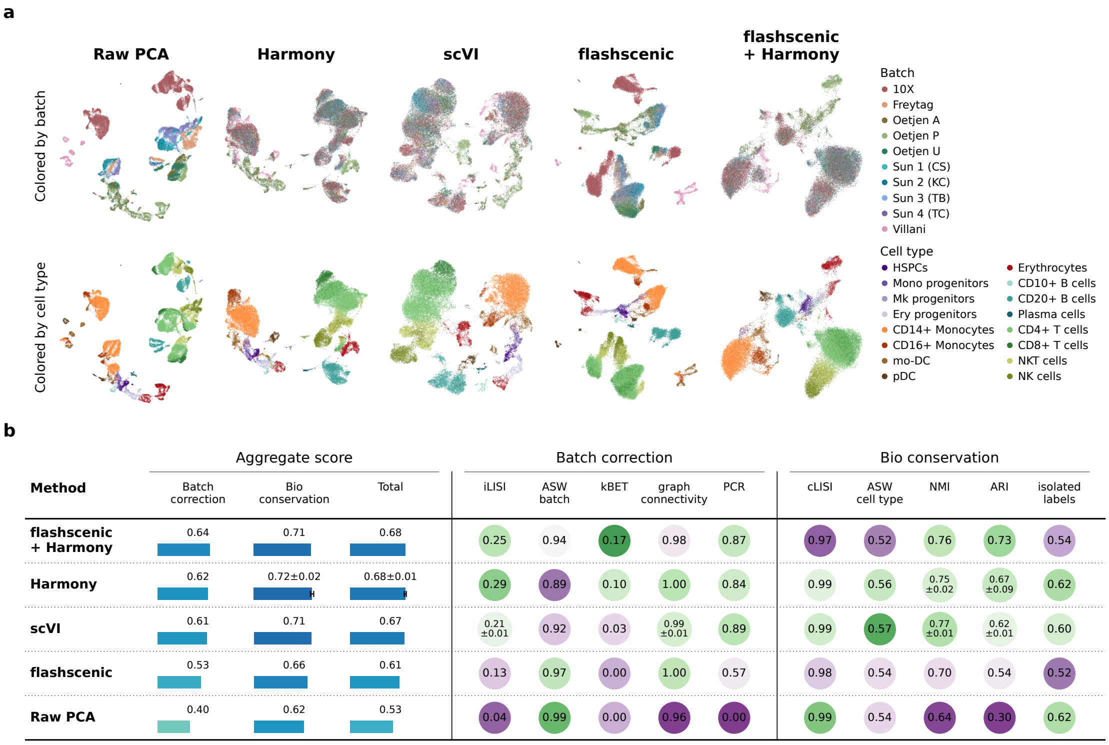


**Fig S4. flashscenic batch integration in the scIB human immune benchmark dataset.** Same layout and methodology as Figure 2b,c (33,506 cells, 10,410 genes, 10 batches, 16 cell types).

**Table S2.** Ranking of the nine consensus aggregation strategies by CollecTRI F1 score (harmonic mean of mean precision and mean recall), averaged across three seed replicates per dataset; in each of the three robustness datasets, and by mean rank across the three. The bottom row gives the random-gene-draw chance-level F1 for reference (not ranked).

| Strategy | F1 (ALS) | F1 (immune) | F1 (lung) | Mean rank |
|---|---|---|---|---|
| Freq ≥50% min5g | 0.0479 | 0.0758 | 0.0742 | 1.00 |
| Freq ≥50% | 0.0426 | 0.0749 | 0.0702 | 2.33 |
| Freq ≥80% min5g (pySCENIC) | 0.0429 | 0.0692 | 0.0648 | 4.33 |
| Freq ≥75% | 0.0400 | 0.0685 | 0.0686 | 5.00 |
| Mean-adj min5g (no CV) | 0.0418 | 0.0633 | 0.0659 | 5.33 |
| Mean-adj + CV<0.25 | 0.0407 | 0.0639 | 0.0656 | 5.67 |
| Mean-adj (no CV) | 0.0405 | 0.0637 | 0.0657 | 6.17 |
| Mean-adj + CV<0.5 | 0.0405 | 0.0637 | 0.0657 | 6.17 |
| Mean-adj + CV<0.1 | 0.0380 | 0.0519 | 0.0508 | 9.00 |
| Expected by chance | 0.0068 | 0.0120 | 0.0114 | — |

Note: Mean rank is the average of each strategy's F1 rank (1 = best of nine) across the three datasets; ties in F1 (e.g. Mean-adj (no CV) and Mean-adj + CV<0.5, whose CV mask removes no edges at this threshold) receive tied ranks. Freq ≥50% min5g ranks first in every individual dataset.

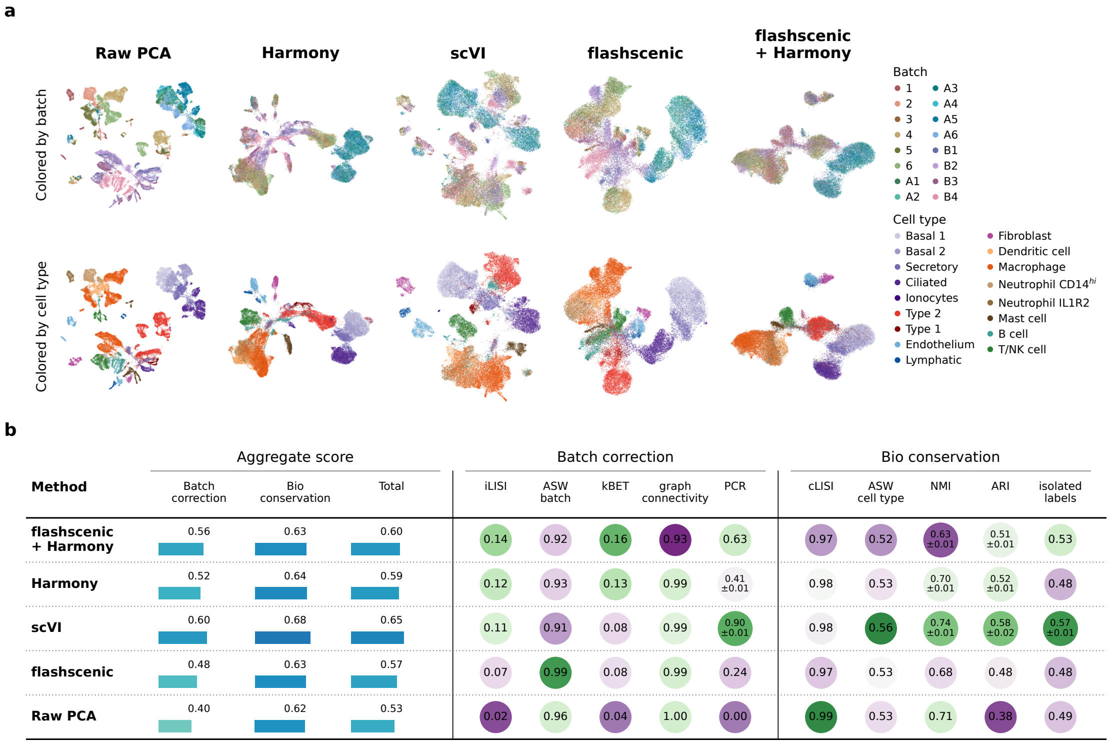


**Fig S5. flashscenic batch integration in the scIB human lung benchmark dataset.** Same layout and methodology as Figure 2b,c (32,281 cells, 13,184 genes, 16 batches, 17 cell types).

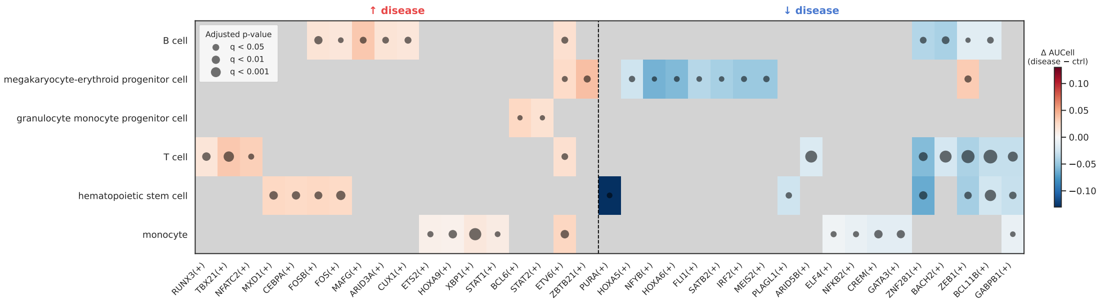


**Fig S6. Differential TF activity across AML bone marrow cell types.** glmmTMB zero-inflated Beta regression (Section 1.5.2) of AUCell regulon activity vs. disease status, donor random intercept, per cell type with $\geq 500$ cells. Top 10 regulons by mean activity difference ($\Delta\overline{\mathrm{AUC}}_{\mathrm{disease-control}}$) per cell type and direction are shown; dot size encodes BH-adjusted $q$-value, dot color encodes $\Delta$ (red: higher in disease, blue: lower in disease).

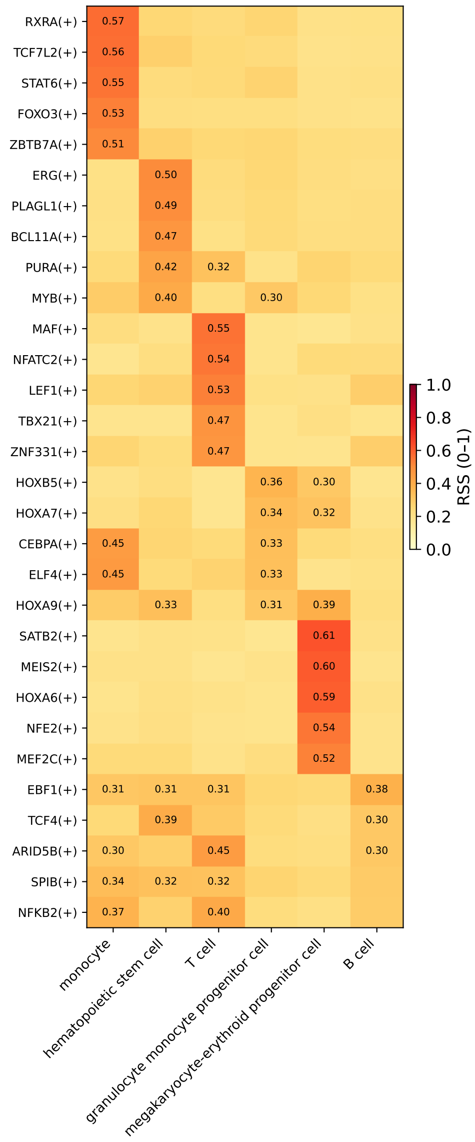


**Fig S7. Cell-type-specific regulon activity in AML bone marrow.** Regulon Specificity Score (RSS) computed on the full 116,043-cell AML dataset, top 5 regulons per cell type, restricted to the six cell types shared with the differential TF activity analysis (Supplementary Fig. S6). Cell values are RSS (0–1); blank cells indicate the regulon is not among that cell type's top 5.

**Table S3.** Pairwise Welch's $t$-test (unequal variance) on Total scIB score among the three seed-replicated methods (5 seeds each; Section 1.5.1), for each of the three datasets. $\Delta$ is method$_1$ minus method$_2$ mean Total score; $q$ is Benjamini–Hochberg corrected within each dataset across its 3 pairwise comparisons.

| | | AML | | Immune | | Lung | |
|---|---|---|---|---|---|---|---|
| Method 1 | Method 2 | $\Delta$ | $q$ | $\Delta$ | $q$ | $\Delta$ | $q$ |
| flashscenic+Harmony | scVI | +0.0290 | $< 0.001^*$ | +0.0111 | $0.001^*$ | −0.0490 | $< 0.001^*$ |
| Harmony | flashscenic+Harmony | −0.0044 | 0.219 | −0.0001 | 0.990 | −0.0096 | $< 0.001^*$ |
| Harmony | scVI | +0.0246 | $< 0.001^*$ | +0.0110 | 0.183 | −0.0586 | $< 0.001^*$ |

*Note:* $^*q < 0.05$.
